# Do Sigmoidal Acquisition Curves Indicate Conformity?

**DOI:** 10.1101/159038

**Authors:** Paul E. Smaldino, Lucy M. Aplin, Damien R. Farine

## Abstract

The potential for behaviours to spread via cultural transmission has profound implications for our understanding of social dynamics and evolution. Several studies have provided empirical evidence that local traditions can be maintained in animal populations via conformist learning (i.e. copying the majority). A conformist bias can be characterized by a sigmoidal relationship between a behavior’s prevalence in the population and an individual’s propensity to adopt that behavior. For this reason, the presence of conformist learning in a population is often inferred from a sigmoidal acquisition curve in which the overall rate of adoption for the behavior is taken as the dependent variable. However, the validity of sigmoidal acquisition curves as evidence for conformist learning has recently been challenged by models suggesting that such curves can arise via alternative learning rules that do not involve conformity. We review these models, and find that the proposed alternative learning mechanisms either rely on faulty or unrealistic assumptions, or apply only in very specific cases. We therefore recommend that sigmoidal acquisition curves continue to be taken as evidence for conformist learning. Our paper also highlights the importance of understanding the generative processes of a model, rather than only focusing solely on the patterns produced. By studying these processes, our analysis suggests that current practices by empiricists have provided robust evidence for conformist transmission in both humans and non-human animals.

## Introduction

There is increasing evidence that many animals, including humans, use social learning strategically; preferentially copying certain individuals, behaviours, or in particular contexts, in order to optimise the quality of information gained^1, 2^. These learning rules will not only determine what behaviours individuals acquire, but will also shape transmission dynamics as information moves through populations, with implications for the formation and patterning of traditions^1, 3–5^. One such social learning strategy is conformist learning, where individuals acquire behaviour with a positive frequency-dependent learning bias, preferentially copying the most common behaviour or the majority of individuals^6, 7^. Conformity has been of particular interest in study of cultural evolution, as it is considered one of the central pillars in the emergence of culture^8^. That is, if individuals copy the majority, groups should fix on a single socially learnt tradition that will remain stable over time and be resistant to erosion or invasion by alternative variants, allowing for group specific cultures to establish and persist. Once thought of as a potentially ‘human only’ trait^6^, numerous models have now argued that conformity could potentially evolve in various species under a wide range of environmental conditions^7, 9^. For example, a conformist learning strategy can be adaptive in spatially varying environments where local knowledge is costly to acquire^3, 8, 10^, as copying the behaviour of the majority of residents will thus be a short-cut to ascertaining best locally adapted behaviour. However, experimentally demonstrating the use of conformist learning in non-human animals has been challenging, particularly in natural populations. A key question has thus become whether conformist learning strategies can be inferred from behavioural data on the acquisition and expression of learnt behaviour.

Recently, two studies provided evidence for conformity in very different systems. van de Waal et al. (2013)^11^ trained troops of vervet monkeys (*Chlorocebus aethiops*) to prefer one of two colours of otherwise identical corn. They found that naÂ¨Ä±ve monkeys adopted the majority behaviour of their group, and that dispersing individuals would switch their choice to match the majority behaviour in the group that they joined. Aplin et al. (2015)^12^ trained great tits (*Parus major*) to access a food reward by solving a puzzle box by sliding a door either left or right, where puzzle boxes could be opened in either direction. They then observed the spread of this new behaviour from these trained demonstrators across five replicated sub-populations and over two generations. NaÂ¨Ä±ve birds disproportionately copied the most common solving technique (opening direction) in their local foraging group, with traditions for left or right persisting across generations. Similarly to the vervet monkeys^11^, birds that moved between sub-populations with different traditions tended to switch their preference to match the locally common behaviour of the area they immigrated into.

These studies reflect the major lines of evidence used to suggest conformist biases in non-human animals. Firstly, that individuals will change their existing behaviour when faced with a conflict between their preferences and those of the rest of their group, and secondly, that the probability of a naÂ¨Ä±ve individual adopting the most common behaviour will be disproportionately greater than that behaviour’s frequency in the group it observes. This second line of evidence can be characterized by a sigmoidal relationship between an individual’s probability of acquiring a behaviour and the frequency of its demonstration. When individuals can choose between two behaviours (sometimes including non-adoption as a behaviour), such a sigmoidal relationship indicates how individuals become disproportionately biased towards the common behaviour^3^. It is important here to differentiate *acquisition curves* from *diffusion curves*, as both are common topics in the study of social learning. A diffusion curve is the instantaneous or cumulative rate of adopting a behavior in population as a function of time^13^. Our focus is on the acquisition curve, which is the instantaneous rate of adopting a behavior as a function of that behavior’s prevalence in the population.

Scaling up, the presence of a sigmoidal acquisition curve at the *population* level—in which the relationship between the frequency of a behavior and the overall rate at which it is adopted is sigmoidal—has generally thought been to provide evidence of conformist learning in the population in question.Yet is this approach robust? Several recent papers have argued that the answer is a firm ‘no’^14–16^. These critiques center on the claim that sigmoidal curves can be generated without conformist learning, and therefore that the presence of such curves is not evidence for conformist learning. In the remainder of this paper, we evaluate these claims.

The evidence for conformist learning in wild populations of non-human primates and birds was first challenged by van Leeuwen et al. (2015)^14^. In this paper, they disputed the methodological validity of using the frequency at which a behaviour is being demonstrated when testing for a conformist bias (for example, as in Aplin et al. 2015a^12^). Instead they argued that a valid test for conformity must compare the probability of adopting behaviours against the overall proportion of individuals performing each behaviour; for example, if a demonstrator performs a behaviour twice, its contribution to the overall frequency of behaviour in a group is only counted once. In response to this critique, Aplin et al. (2015b)^17^ reanalysed their previous data to show that their results were robust to either definition of conformist transmission. Aplin et al.^17^ further argued that, in empirical practice, there were no cases in which the two definitions conflicted, suggesting that they are functionally equivalent in all but the most exceptional of cases^17^. This argument was further supported by re-analyses of data from van de Waal et al. (2013) by Whiten and colleagues^18^.

In two more recent papers, van Leeuwen et al. (2016)^15^ and Acerbi et al. (2016)^16^ have issued a more direct challenge to the validity of identifying conformist transmission from population-level sigmoidal acquisition curves. In these papers, they developed a simple computational model in which populations of 100 individuals were randomly initialized with one of two behaviours. Individuals were then allowed to either copy a random demonstrator^15^ or choose to copy a demonstrator as determined by a range of different learning rules^16^. In several cases, their modelling results suggested that plotting the frequency of the behaviour against the probability of adoption could lead to sigmoidal acquisition curves even in the absence of conformist learning. In particular, Acerbi et al. (2016)^16^ purported to demonstrate two non-conformist learning rules that nevertheless generate conformist curves: one in which individuals have an intrinsic bias for one behaviour (â€˜variant preference’), another in which individuals copy a behaviour from a randomly chosen member of a small subset of the population (’demonstrators subgroup’).

Here, we revisit the modelling approaches used by van Leeuwen et al (2015, 2016)^14, 15^ and Acerbi et al (2016)^16^. Specifically, we reanalyse the models in two regards. First, we examine the evidence that proposed non-conformist learning mechanisms can yield sigmoidal curves. We then consider the argument that the majority should be represented as the frequency of individuals exhibiting a behaviour, rather than by the frequency of the behaviour itself. This reanalysis highlights that in both cases, a number of problematic assumptions and implementation errors distort or exaggerate the results produced. Our aim in this paper is therefore to correct the overly strong claims made in those papers. We conclude that, with proper methodological implementation, a sigmoidal relationship between the adoption of a behaviour and its frequency in the group remains a valid and robust way to identify conformist social learning in wild populations.

## Results

### Do non-conformist social learning strategies generate sigmoidal curves?

Acerbi et al. (2016)^16^ used an individual-based model to investigate when a sigmoidal relationship arises between the proportional frequency of a behavioural variant A and the probability of that a random individual will adopt A over a competing variant B. The authors examined 10 possible learning biases, and concluded that two non-conformist learning biases could also produce sigmoidal curves: ‘variant preferences’, and ‘demonstrator subgroups’. We address each of these in turn (see also Table 1 for a summary). Before doing so, we first address an underlying assumption of their model that has important implications for any interpretation of its results: the 50/50 variant starting condition.

**Table 1.** Summary of problematic issues found in the models by Acerbi et al (2016) and Van Leeuwin et al (2016). These issues result in the incorrect or misleading conclusions drawn from the simulations. While some issues do not strongly affect the shapes of the curve in the results of the papers, they incorrectly implement the biological process that the papers seek to investigate (for example, animals do not copy themselves) or incorrectly represent the analytical approaches used by empirical studies (for example studies count only the first demonstration by an individual).

| Issue | Explanation |
| --- | --- |
| Starting conditions | The population was always initiated with a probability of 0.5 for each behaviour. This is unrealistic from empirical perspective, as a natural spreading process would start with one innovator (proportions near 0 or 1). This assumption exacerbates the potential for an artefactual sigmoidal response. Had range of initial conditions been used to assess model robustness, which is common practice in agent-based modelling, the strong claims made by the modelling papers would not have been supported. |
| Pooling data | All of the frequency data across each of the 1000 replicates were pooled, without noting variation across replicates. This aggregates multiple categories of response into a single category. This practice can eliminate discovery of multimodal data and provides no information on the range of outcomes that are possible. This approach can graft different responses together, such as combining two linear responses that then appear as an artefactual sigmoid. More generally, data with categorically different starting conditions should not be pooled. |
| Full behaviour history | The full history of all behaviours was used to calculate frequency of behaviours used by agents in the model when making a behavioural choices. This approach misrepresents what has been done in existing empirical studies. It also describes a biologically implausible process that assumes individuals respond to a distant past that potentially includes thousands of observations. Using data restricted to recent history when calculating the state of the system alleviates any potential mistaken results. |
| What counts as ‘copying’ events | Every event was counted in which an individual encountered behavioural variant A, including when an individual did not change what action it performs. However, events in which an individual *retained* variant A after encountering variant B were not counted. This increases the appearance of a sigmoidal acquisition curve (as opposed to either a linear or r-shaped curve) because individuals that continuously perform variant A when the population behaviour becomes entrenched are all counted as *copying* action A. Defining social learning events is challenging, so striving for consistency and robustness is important. |
| Individuals are their own targets for social learning | Individuals could select themselves as demonstrators, which confounds data on social learning. Similarly, a focal individual’s behaviours were counted in its assessment of the historical frequency of observed events. Whilst in a population of 100 or more individuals this issue had relatively little impact on the results, the effect may be much greater in a smaller population. |

#### Starting conditions

In the model of Acerbi et al.^16^, a population of 100 individuals is initialized in which approximately 50% adopt behavioural variant A, and the rest adopt variant B. That is, the probability that an individual performs behaviour A at time = 0 is, on average, 0.5. They then simulate 10000 copying events in which individuals randomly copy the behaviour of another individual, and use a range of rules for selecting these individuals. They record each of these copying events as the history of events. They replicate this process 1000 times. Data is then aggregated across all model runs for the entire length of each simulation in order to describe the full range of variant frequencies, as either A or B goes to fixation. At least two potential problems arise from this methodology. First, in the absence of group structure or anti-conformist drives, a random copying rule implies that the absorption probability increases with the frequency of the variant; in expectation, the population monotonically converges to a single dominant variant. Thus, data for frequencies of variant A between 0.5 and 1 will come exclusively from runs in which uniform adoption of variant A was the absorbing state, while data for frequencies of variant A between 0 and 0.5 will come from runs in which the absorbing state was variant B (with some exceptions for stochastic noise around frequencies of 0.5). A corollary is that the dynamic in which a rare variant arises and becomes dominant is never explored. Second, because data is aggregated from across all time steps, more common variant frequencies will be overrepresented in the data. This should not matter if all aggregated populations are drawn from the same distributions. However, as we show below, errors can arise when populations using disparate rules are averaged together.

#### Variant preference

Acerbi et al. (2016)^16^ claim that a non-conformist copying rule, which they call variant preference, can generate sigmoidal curves. The rule is instantiated in the model as follows:

> Individuals had a preference for one of the two variants, which was operationalized by endowing individuals with different copying probabilities for the two variants. When presented with a demonstrator showing it, one of the variants was always copied, while the other variant (the less preferred) was copied according to the parameter *pLess* (*pLess* was varied between 0 and 1). All individuals preferred the same variant, which was randomly selected at the beginning of each repetition. (Acerbi et al. 2016, p. 7)^16^.

The sigmoidal curves generated stem from the two faulty assumptions described above: averaging data across all runs, and always starting with a 50-50 mix of behavioural variants^16^. We first replicate their result (Fig. 1, top row middle) of a sigmoidal curve. We then separate runs in which variant A is preferred (Fig. 1, top row right) from runs in which variant B is preferred (Fig. 1, top row left). This indicates that the sigmoidal curve is actually an artefact formed by joining together two lines with different slopes. What explains this?

**Figure 1.**
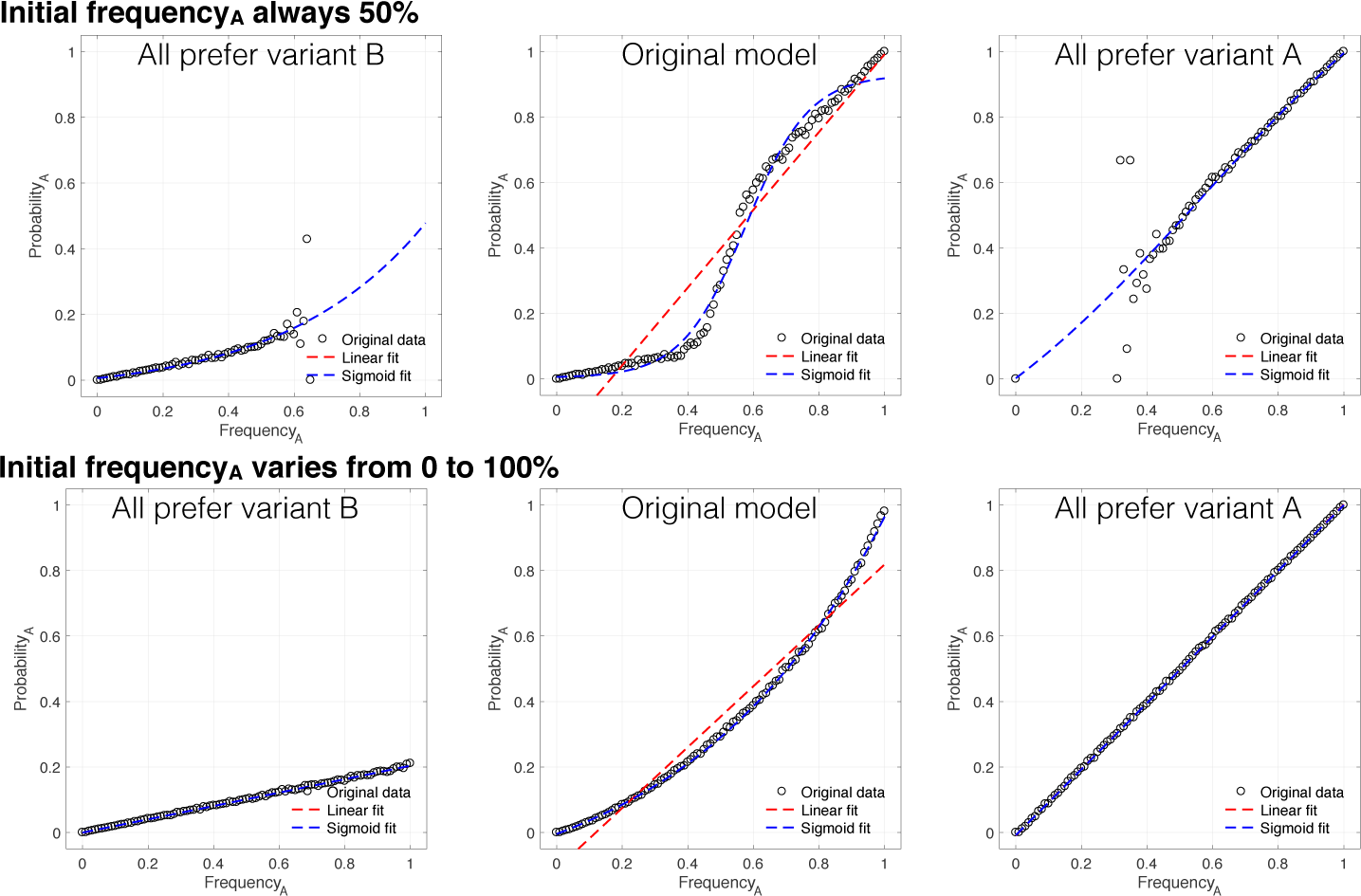
Variant preferences. A replication of the model of Acerbi et al.^16^ in which individuals prefer one of the two variants. Includes best fit linear and sigmoid curves to data. Top row: Initial frequency of variant A always initialized to 50%, as in Acerbi et al. Bottom row: Initial frequency varied across runs from 0% to 100% in increments of 0.1%. Centre column: Runs in which variant A is preferred and runs in which variant B is preferred are averaged together. Right and left columns: variant A or B is always preferred, respectively.

When each individual prefers variant A, the probability of adopting A is simply equal to the probability of encountering an individual exhibiting variant A, which is just the frequency of A. For each individual preferring variant B, the probability of adopting A is the frequency of A times *pLess*: the probability of encountering an A times the probability that A is adopted in those cases. The sigmoidal curve results from these two lines being stuck together. Of course, two lines being averaged together should not result in a sigmoidal curve. This feature stems from the fact that the frequency of each variant is always initialized at 50%. Because adoption dynamics are always monotonic in expectation (i.e., the frequency of one variant always steadily increases, stochastic noise notwithstanding), the left and right sides of the graphs are generally from completely different batches of simulations, as illustrated by the top row of Fig. 1. In most empirical studies, individuals in the population are not initiated with a particular behaviour at random, but rather will start out naÏve, so that the probability of encountering a behavior will start close to 100%, rather than 50%. We ran simulations with a uniform distribution of initial adoption frequencies for variant A ranging from 0% to 100%. The results (Fig. 1, bottom row) illustrate how preferences for either variant are associated with a straight line with slope depending on preference strength. Combining data from both preferences reveals a concave-up curve rather than sigmoidal curve. Such a curve, rather than a straight line, is a result of more data for the dominant variant preference at either end of the frequency spectrum. A simple test supports this interpretation (Fig. 2).

**Figure 2.**
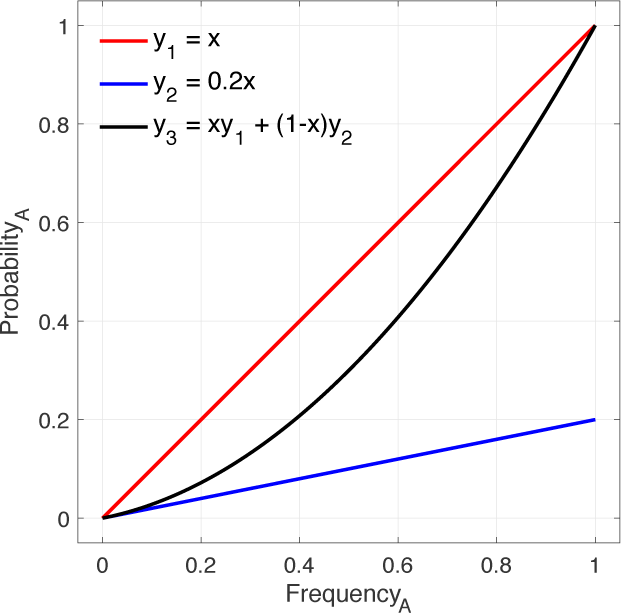
Explanation of concave-up curve. Curves *y*_1_ and *y*_2_ are simple lines with slopes of 1 and 0.2, reflecting the probability of adopting variant A or B, respectively. Curve *y*_3_ is the weighted average of the two lines, in which *y*_2_ for lower values of *x* and *y*_1_ is dominant for higher values. Note that *y*_3_ replicates the curve seen in Fig. 1, bottom centre.

#### An analytical model of variant preference

The variant preference rule is equivalent to what Boyd and Richerson (1985)^3^ called a direct bias. It is easy to derive a simple mathematical model of such a bias, and doing so will help to illustrate the dynamics of the individual-based model discussed above. Each individual observes another individual at random and decides whether to adopt their behaviour. Let *p* be the frequency of variant A. Upon observation, the probability of adopting one’s preferred variant is 1, and the probability of adopting the less-preferred variant is *q* (equivalent to *pLess* in the Acerbi et al. model^16^). Finally, let *r* be the frequency of individuals who prefer variant A, in order to accommodate a mixed population of preference types. It now follows that the probability that a randomly chosen individual adopts variant A is given by:

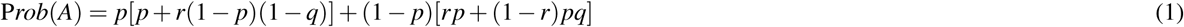

The first term on the right side of this equation is the probability that the observer initially exhibits variant A and continues to do so after the observation, while the second term is the probability that the observer initially exhibits variant B but switches behavior after the observation.

Focusing on variant A, we see that if everyone in the population favours the same variant, we should observe strict concave up or down curves, as in Fig. 3A. If there is an even mix of both types, or if we average over an even number of runs of both types, the curves will average out, and we end up with a straight line. However, a sigmoidal curve can once again be generated from the assumption that the population is always initialized with 50% of each variant. Whenever one variant is favoured, that variant will always increase, and so no data will ever be recorded for frequencies less than 0.5. Rather, frequencies below 0.5 of variant A will only be observed when variant B is preferred (r < 0.5). Combining these two situations leads to the union of two distinct curves, and can generate an artefactual sigmoid curve, as in Fig. 3B. However, this bears little resemblance to any realistic data that would be collected in an empirical study.

**Figure 3.**
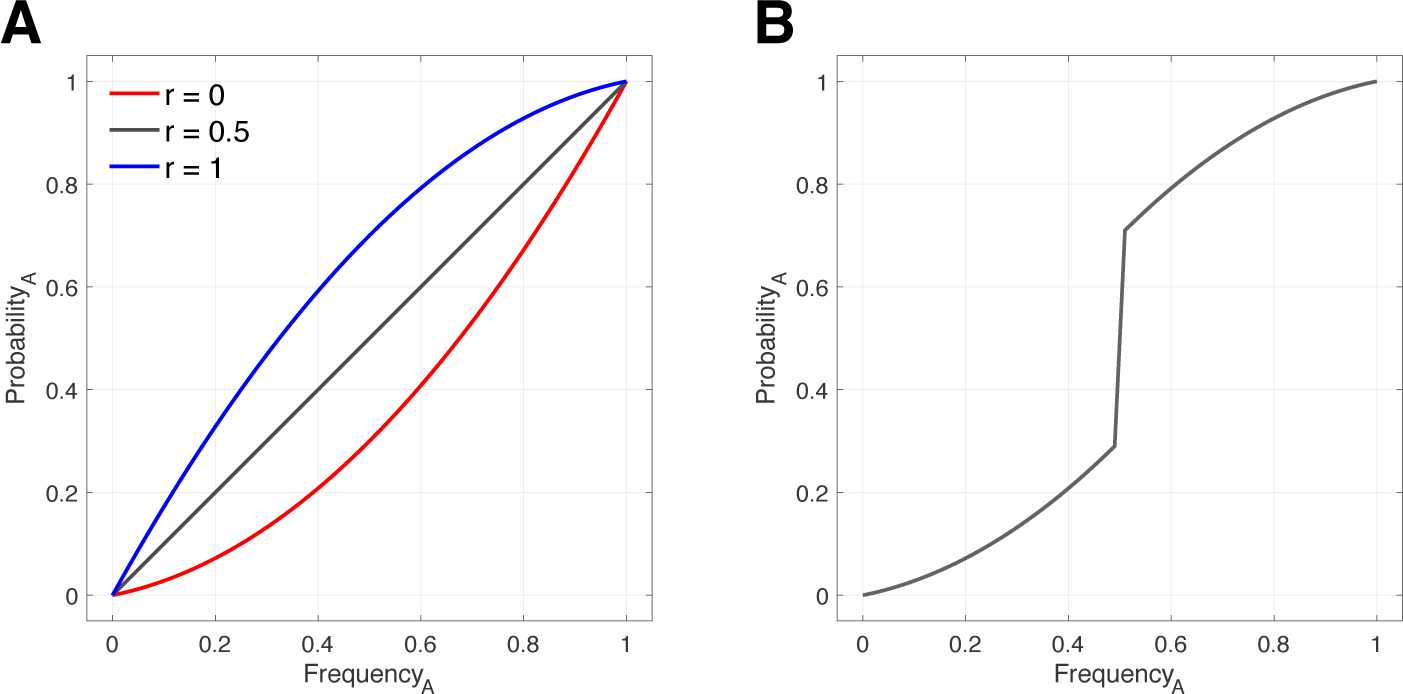
Analytical model for variant preferences. (A) When all individuals prefer variant A (*r* = 1), we replicate the r-shaped curve of Boyd and Richerson (1985)^3^. (B) If the environment is always initialized with approximately 50% of each variant and the preferred variant always increases in frequency, an artefactual sigmoidal curve can emerge if all data for both variant preferences are combined. To be consistent with agent-based simulations, curves here are for *q* = 0.2.

Why does this analytical model give an r-shaped curve for the case where all individuals prefer variant A, while the model of Acerbi et al.^16^ gives a straight line? The difference allows us to notice yet another important modelling artefact. Acerbi et al. took as their dependent measure the probability that any randomly selected copying event in which the focal individual was exposed to another individual with variant A resulted in that focal individual adopting variant A. This counts events in which the focal individual was already exhibiting variant A and then encountered an A, and so “copied” the variant she already had. However, it does *not* count events in which the focal individual started with variant A and encountered a B, but refrained from copying the newly encountered variant.

In our analytical model (Fig. 3), the dependent variable is, instead, the probability that a random individual will end up demonstrating variant A, given the current frequency of individuals exhibiting variant A in the population. In other words, the analytical model looks at the behaviours expressed in the population regardless of who encountered whom, and so counts the case where a focal individual started with variant A, encountered a B, and refrained from copying the newly encountered variant. Thus, the probability of demonstrating variant A when it is preferred is higher in our model than in the model of Acerbi et al.^16^. We view this as a more reasonable assumption, since the probability of adopting a behavioural variant should be enhanced by exhibiting an internal bias for that particular variant. We note also that our analytical result mirrors a very similar result obtained previously by Boyd and Richerson (1985, p. 207)^3^.

We also believe that the dependent measure of adoption used in our analytical model is more in line with empirical methodology than the measure of adoption used by Acerbi et al.^16^. The sample sizes that empirical studies deal with will often be substantially smaller than the size of the populations being modelled. A consequence of this is that researchers will use all of the data available to them (i.e. not ignore some cases—as when individuals starting with variant A encounter B and refrains from copying it).

In summary, the variant preference copying rule does not generate sigmoidal curves when unrealistic modelling assumptions are removed. By investigating *why* a sigmoid was produced in the model by Acerbi et al.^16^, we have shown that the variant preference rule is not a valid competing hypothesis for conformist learning when considering the causes of sigmoidal acquisition curves in results from empirical studies.

#### Demonstrator subgroups

The second copying rule purported by Acerbi et al. (2016)^16^ to yield sigmoidal acquisition curves was called demonstrators subgroup. This rule corresponds to a situation in which individuals exhibit a model bias, for example by copying dominant or prestigious individuals (e.g., a ‘copy dominant males’ rule), and works as follows:

> At the beginning of each repetition, a subset *S* of individuals in the population, of size *Dm* (*Dm* = 5, 10, 20, 50), was randomly chosen. At each time step, each individual was paired with a demonstrator chosen from the subset *S* and always copied it. Demonstrators themselves changed their state according to the same rule. (Acerbi et al., p. 7)^16^.

We first replicate the results of Acerbi et al.^16^ (Fig. 4A). Unlike variant preference, the sigmoidal curve generated is robust to variation in the initial frequency of each variant when averaged across simulations using a uniform distribution of initial frequencies (Fig. 4B). If, however, the behavioural variant that is preferentially copied is initially rare, the sigmoidal curve becomes increasingly skewed, approaching an r-shaped curve (Fig. 4C). The reason the ‘demonstrators subgroup’ rule generates a sigmoidal curve is as follows. Due to the well-known phenomenon of drift in small groups^19, 20^, random copying among the demonstrators will soon create a majority and then a uniform bloc. When one variant has a majority in the demonstrators subgroup, that variant will be copied at a rate higher than chance, because its frequency will increase more rapidly in the small subgroup than in the population at large. Once the variant reaches fixation in the subgroup, it will be copied with certainty by members of the population, evidenced by the horizontal flattening out of the acquisition curve for small and large frequencies of variant A in Fig. 4A (the severity of these lines is reduced in Fig. 4B because the initial frequencies of the variant in the subgroup and in the larger population are less strongly coupled). By restricting analysis to conditions in which variant A is initially rare, we lose the strong horizontal line for low frequencies of A (Fig. 4C), in part because adopting variant A will in those cases be a rare event.

**Figure 4.**
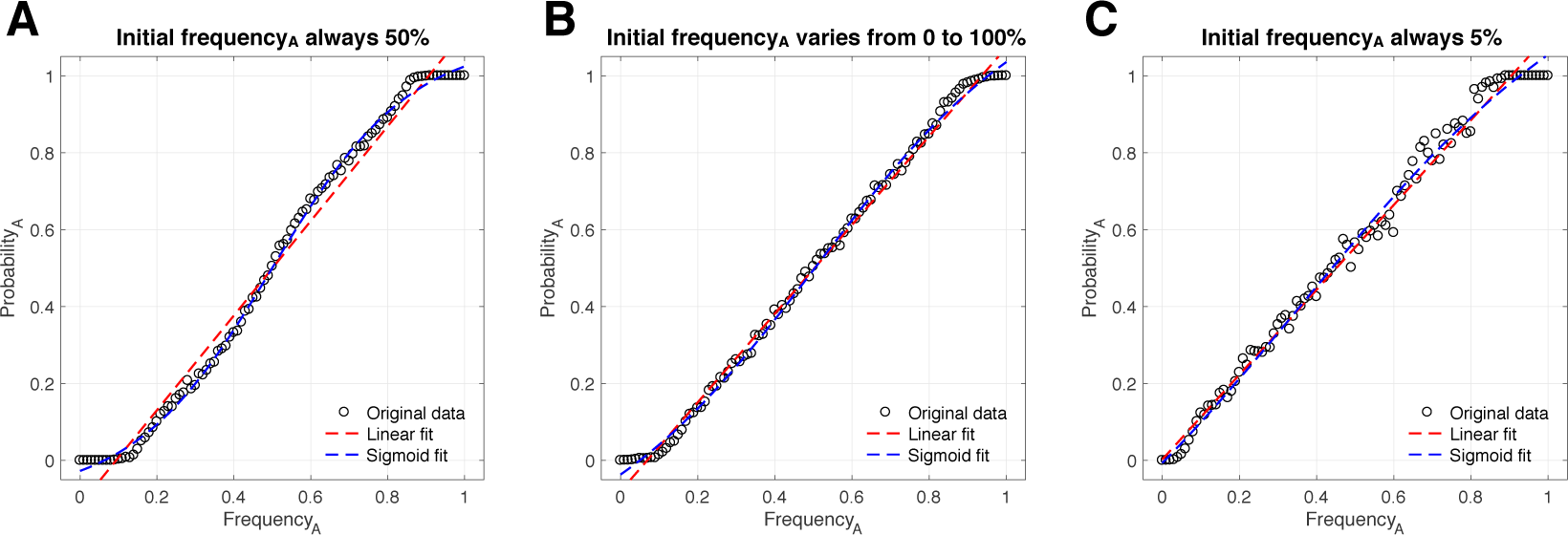
Demonstrators subgroup. Individuals choose a random member of a fixed group of size *Dm* and copy that individual’s behavioural variant. For all simulations, *Dm* = 5. (A) Initial frequency of variant A is fixed at 50% for all runs, replicating the results of Acerbi et al. (2016)^16^. (B) The sigmoidal curve remains apparent when initial frequency of variant A is varied across runs from 0% to 100% in increments of 0.1%. (C) When variant A is always rare initially, the sigmoidal curve largely disappears, replaced with an r-shaped curve.

The acquisition curve generated by the ‘demonstrators subgroup’ rule is more severe (characterized by larger jumps in the second derivative) than the curve generated by conformist transmission; however, it can be classified as a sigmoidal curve. Should researchers therefore be concerned that their sigmoidal data may reflect a model bias (e.g. a strategy of copying dominant individuals) rather than a majority bias? Probably not. Acerbi et al. (2016)^16^ showed that the demonstrators subgroup rule reliably generated a sigmoidal curve only when both the total group size and the relative number of demonstrators were very small. Specifically, a sigmoidal curve provided a better fit than a line only when *N* ≤ 100 and *Dm* < = 5. Practically speaking, this is likely equate to exclusively copying one or a few specific individuals, rather than preferentially copying a type or class of individual. In such cases, it should be relatively easy for researchers to identify this sort of dynamic and to test the demonstrators subgroup hypothesis against more direct conformist biases absent dominant leaders.

### How to measure the majority

Conformist learning occurs when the more common behavioural variant is preferentially adopted. Empirically, there are at least two ways to measure just *how* common a particular variant is. You can count the number of individuals exhibiting each variant, or you can count the total instances of each variant. In the latter case, some individuals may be counted more than once. Two papers by van Leeuwen et al.^14, 15^ purported to show how using the frequency of observed behaviours as a measure of the majority, rather than the frequency of individuals exhibiting those behaviours, can lead to incorrect conclusions about the presence of conformist learning. To illustrate this, van Leeuwen et al.^15^ constructed a model similar to the above, and compared acquisition curves produced using the frequency of behaviours and the frequency of individuals. They found that measuring the frequency of behaviours could produce artefactual sigmoidal curves even when individuals copied the behaviors of randomly chosen individuals, and so claimed that a reliable indicator of conformity could be derived only by measuring the frequency of individuals.

Firstly, it is worth noting that when implementing their model, the authors recorded the complete history of behaviours expressed by the population up to the point where the observer made their decision. This implementation suggests that observers use the complete history of all behaviours exhibited by all individuals when choosing between variants, or at least that empirical researchers analyse their data under this assumption. We would argue that this is highly implausible; rather, copying individuals will observe only a small subset of the population during a decision-making event. Ideally, researchers would like to be able to quantify exactly what focal individuals have actually paid attention to prior to exhibiting their behaviour. To date, this has been methodologically challenging, and so instead most researchers have recorded the frequency of behaviours (or of individuals undertaking behaviours) in some constrained time prior to the focal individual making its decision. This approach attempts to estimate what the individual might have plausibly been able to observe in its current learning period. For example, Aplin et al. (2015)^12^ used a 245 second period prior to each bird’s first solve (a mean of 6 previous solves) as their ‘history of behaviour’ time-window, and based this value on behavioural patterns observed from extensive studies in their population.

Here we re-ran the baseline model from van Leeuwen et al (2016)^15^ and Acerbi et al (2016)^16^, in which individuals randomly select one individual to copy (Fig. 5A). Taking the entire history of behaviours produced by individuals in the population replicated the same sigmoidal response that was produced in the original studies (Fig. 5B). However, if the history was limited to the previous 10 behaviours, we found that the relationship was perfectly linear, as expected under a random copying strategy (Fig. 5C).

**Figure 5.**
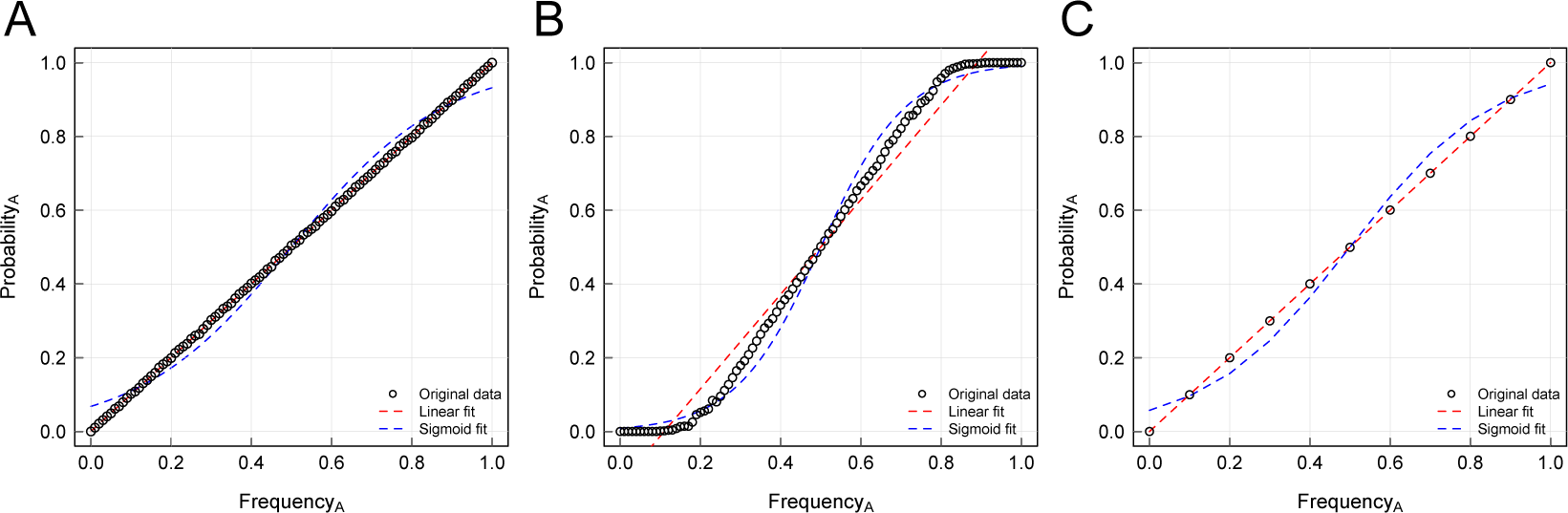
Implementations of the model described in van Leeuwen et al., counting the behavioural variants from the current state of the population (A), and from the complete history of actions performed (B). If the history of actions performed is limited to a recent window (here the last 10 solves, (C)), then there is no evidence that a sigmoidal curve could result.

However if one is not going to use the complete history of behaviours, then how long should the time window for data collection be? A limitation of using a time window approach for considering the behavioural history of demonstrations is that the size of the time window is not always obvious, as highlighted above and in the critique by van Leeuwen et al (2016)^15^. Could the choice of time-window influence the likelihood of erroneously producing a sigmoidal curve in the absence of conformist copy? To test this, we measured the sensitivity of the relationship between frequency of behaviours observed and the probability of performing the behaviour to the size of the time window (Fig. 6). We found that a linear model provided a much better fit to the data than a sigmoidal model up to window sizes that were still quite large (more than 2000 observations). From this, we can conclude that an incorrect inference of conformist transmission when copying is in fact random will be very unlikely if any time-windowing is applied when calculating frequency of behaviours.

**Figure 6.**
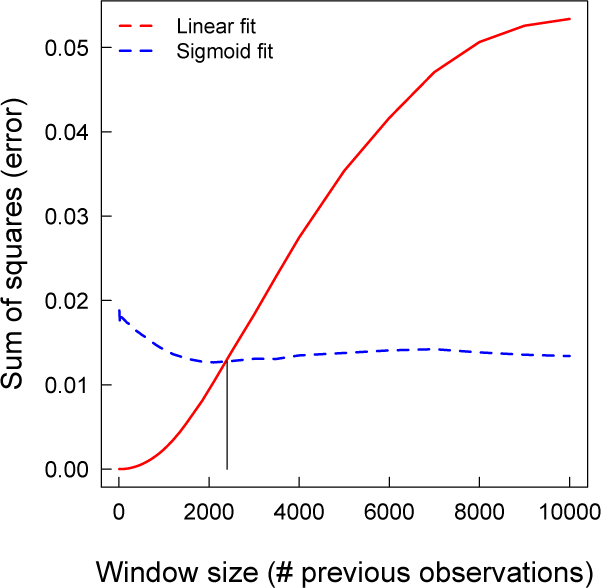
Relative fit of a linear versus sigmoidal model based on the *r*^2^ value of the model fit. Here a window of size *x* is used to calculate the variant frequencies, where *x* is related to the number of previous observations from the population. As *x* increased, the fit of a linear model (solid line) decreases. However, a linear model remains a better fit than a sigmoid model until a very large window size.

## Discussion

Three recent studies have questioned the empirical evidence for conformist copying in non-human animals, and re-opened the debate as to whether sigmoidal acquisition curves can be used as evidence for conformity. In this paper, we examined the different processes that generated sigmoidal curves in these published models, and find that non-conformist learning processes for sigmoidal outcomes were almost exclusively underpinned by flawed or unrealistic modelling assumptions. Our work further highlights how these assumptions misrepresent the types of analyses conducted by empiricists. More specifically, we first examined the non-conformist learning mechanisms purported to yield sigmoidal curves. We showed that one of them, variant preference, failed to do so when more realistic behavioural and analytical assumptions were used. The second mechanism, demonstrator subgroup, can generate weakly sigmoidal curves under some conditions, but is unlikely to be mistaken for conformist transmission and is easy to control for empirically. Second, we considered the argument that the majority should be represented as the frequency of individuals exhibiting a behaviour, rather than by the frequency of the behaviour itself, as the latter can also generate sigmoidal patterns under random copying. We showed that this conclusion stems from an unrealistic modelling assumption that bears little relevance to the overwhelming majority of empirical research.

Our motivation for this paper is neither to provide a general critique of an individual-based modelling approach for understanding social learning and animal behaviour, nor to critique the general caution that alternative generative mechanisms at the individual level may be responsible for population-level patterns. Formal models, including individual-based models, have been critically important in the development of rich theories for understanding social behaviour and evolution. We are also sympathetic to the view that there can exist multiple causal pathways for population-level phenomenal; see, for example, Frank (2009)^21^ on why power laws appear ubiquitous in complex systems. Rather we support publishing guidelines highlighting common errors, in line with the current movement towards a more robust scientific literature^22–25^. Indeed, our study highlights the importance of carefully exploring the factors that drive patterns in a model. A mathematical or computational model acts as a sort of logical engine for turning assumptions into conclusions^26, 27^. Those assumptions concern the delineation of the parts of a system (often individual organisms), the properties and initial distributions of those parts, and the dynamic interactions by which the population changes^28^. The validity of the conclusions drawn from a model therefore depends fundamentally on the extent to which its assumptions adequately represent reality. All models must make unrealistic assumptions if they are to act as useful abstractions. However, the robustness of any model’s results to idiosyncratic assumptions must be carefully assessed if we wish to avoid drawing erroneous conclusions when constructing analogies between models and reality.

Individuals can exhibit various attentional biases during the process of social learning that will have consequences both for acquisition patterns and for what traditions become established at the group level^1, 2, 29^. While a conformist learning bias is one the clearest examples of such an effect, multiple different social learning strategies have been identified across a wide variety of species, many of which could potentially also influence transmission dynamics^1, 4, 13^. It is therefore a valuable exercise to explore whether different social learning strategies could produce similar acquisition patterns, and we applaud the effort to do so. In our reanalysis of Acerbi et al.^16^, we conclude that, indeed, one such social learning rule *can* generate sigmoidal acquisition curves superficially similar to conformist learning: the exclusive copying of a subgroup of potential demonstrators, a form of model bias here called the ‘demonstrator subgroup’ rule. However, it is important to note that this rule only generates a sigmoidal curve when both total group size and the number of demonstrators are small—that is, for populations of 100 individuals or fewer, with five or fewer demonstrators—and even then the sigmoidal nature of the curve is apparent only for very low and very high variant frequencies. For example, a social learning biases such as ‘copy adult males’ would be very unlikely to generate such a acquisition pattern, while ‘copy the single most dominant male’ might. This rule is therefore important to consider as an alternative explanation. However, we believe that empirical researchers have shown themselves to be capable of identifying candidate social learning strategies that might be operating in their system, and devising experiments to test hypotheses against one another. Two recent examples include movement direction in baboons^30^, for which conformist learning was supported, and nest site navigation in honeybees (reviewed in Seeley 2010^31^), in which the preferentially copied individuals were readily identifiable. Finally, ‘copy the dominant’ biases have been explored in vervet monkeys^32^ and thoroughly discussed in the context of additional evidence for conformity in this species^11, 18^.

Does a sigmoidal acquisition curve at the population-level uniquely indicate conformist transmission? We cannot say for certain. However, the recent works by van Leeuwen et al.^15^ and Acerbi et al.^16^ provide no credible evidence that alternative mechanisms could generate such curves under reasonable conditions. This does not mean that no such mechanism exists, nor does it eliminate the possibility that environmental or network constraints could contribute to the emergence of sigmoidal acquisition curves in some circumstances. Indeed, it remains an important question as to whether alternative mechanisms can produce those stable and resilient group-level cultures that are so indicative of conformity^3^, and we encourage further research on this topic. However, if such conditions exist, they are not any that have been previously proposed for the social acquisition of information. In the end, a population-level acquisition curve is a somewhat crude tool with which to investigate the cognitive and behavioural strategies used by individuals. More complex methods and models that can incorporate things like individual heterogeneity, population structure, and the possibility that individuals use multiple learning strategies simultaneously are surely needed^33–35^. Yet for the time being, in the presence of a sigmoidal acquisition curve, conformist transmission should be viewed as a highly probable antecedent.

## Methods

In this study, we re-evaluated the modelling results described in Acerbi et al.^16^ and Van Leeuwen et al.^15^. Our implementations relied on the MATLAB source code provided by Acerbi et al.^36^. We modified this code in only two ways. The first modification was to relax the starting conditions in which the probability for each variant was 0.5 (Figures 1, 4). The second modification is to record, after each copying event, the frequency of behaviours over the previous *T* solves, where *T* = 10 (Figure 5) or ranged from 0 to 10000 (Figure 6). In Table 1, we list some additional implementation issues that we identified while working these models. However, because we were interested in isolating and demonstrating the primary mechanisms underpinning the apparent emergence of sigmoidal curves under random copying, and we did not otherwise alter the original code provided by the authors. Following Acerbi et al., all acquisition data were plotted along with their best-fit linear and sigmoid curves. Analysis of the time window size was based on a replication of the model of Van Leeuwen et al.^15^ in R, the code for which is provided in the online supporting materials.

## Acknowledgements

We are grateful to Alberto Acerbi and Edwin van Leeuwen for their gracious correspondence, and also for sharing their code, without which our analysis would have been considerably more difficult. We thank Brendan Barrett, William Baum, Bret Beheim, Karl Frost, Richard McElreath, Matt Miller, Tom Morgan, and reading groups at UC Davis and MPI-EVA for helpful feedback on an earlier version of this manuscript. L. M. A. was supported by a Junior Research Fellowship at St John’s College, University of Oxford, and both L.M.A. and D.R.F. were supported by a grant from the Biotechnology and Biosciences Research Council (BB/L006081/1).

## Author contributions statement

All authors conceived of the project and helped write the manuscript. P.E.S. and D.R.F. performed the analyses. All authors reviewed the manuscript.

## Additional information

None of the authors have any competing financial interests.

